# Quantifying Assay Confidence in Barrier Organ-on-Chip Systems: A Monte Carlo Assay-Readiness Framework for Detecting Modest Permeability Shifts

**DOI:** 10.64898/2026.09.26.754630

**Authors:** Paola Dama, Shrawasti Sahare

## Abstract

Organ-on-chip (OoC) systems are increasingly used to generate human-relevant evidence in drug discovery and preclinical development, yet interpretation can be limited by the interaction of biological variability, device-to-device variability, and technical assay noise. Here, we describe the UstarFlowAI barrier-confidence framework, an in silico assay-readiness approach that quantifies whether a predefined permeability-related effect is resolvable under specified uncertainty. The proposed context of use is decision support for laminar, low-Reynolds-number airway epithelial monolayer barrier OoC assays in which device fabrication, pump performance, flow, wall shear stress, and biological fluctuation may affect permeability-related readouts. Monte Carlo simulations propagate defined manufacturing, biological, systemic, and well-level uncertainty and estimate the minimum detectable shift (MDS) under a prespecified decision rule. In the present proof-of-concept design, the reference workflow was parameterized at 9.4% technical coefficient of variation (CV), whereas the standardized workflow was parameterized at 5.0% CV, corresponding to an approximately 46.8% reduction in the modeled technical-noise component. At n = 48 replicate wells per group and a nominal total assay-noise operating point of 17% CV, a predefined 10% permeability shift exceeded the upper MDS uncertainty bound for the standardized workflow in both best- and worst-case scenarios, whereas it overlapped the 95% MDS interval for the reference workflow and therefore did not meet the prespecified PASS criterion. These findings demonstrate the behavior of the computational framework under the stated assumptions; they do not constitute experimental validation, regulatory qualification, or evidence that a 10% shift is universally meaningful across barrier OoC systems. We therefore propose a prospective wet-lab validation strategy in which model predictions are tested against airway barrier measurements generated under controlled sources of technical and biological variability. The framework is intended to complement, rather than replace, the underlying biological model by making assay uncertainty and decision thresholds explicit and auditable.

## 1. Introduction

Organ-on-chip (OoC) platforms are increasingly used to model human tissue barriers, including the airway epithelium, where microfluidic systems can support differentiated epithelial cultures and quantitative barrier measurements [1–4]. Flow and wall shear stress are biologically relevant variables in airway models and can alter epithelial barrier function and permeability-related phenotypes [5,6]. These features make barrier OoC systems attractive for mechanistic and translational studies, but they also introduce engineering and assay variables that must be characterized when small treatment-related effects are used for decision-making.

The scientific utility of an OoC assay therefore depends not only on biological relevance but also on the confidence with which a change of interest can be distinguished from experimental uncertainty. Barrier readouts such as apparent permeability, transepithelial electrical resistance (TEER), cytokines, and other functional measurements can be affected by device geometry, flow conditions, manufacturing variation, batch effects, operator or protocol effects, and biological heterogeneity [3,7,8]. More broadly, quality-management and interlaboratory studies in microphysiological systems (MPS) have emphasized the importance of fit-for-purpose performance criteria, reproducibility, and explicit characterization of variability [9,10].

Despite this progress, there is no single, widely adopted quantitative framework for translating device-, flow-, and assay-level variability into a decision boundary for a predefined permeability effect. In particular, technical noise arising from fabrication, pump operation, and assay execution is not always propagated in a way that directly informs replicate design or interpretation of modest treatment-control differences. This gap can make it difficult to determine whether a nominally small effect is genuinely resolvable under the intended operating conditions.

The present work introduces the UstarFlowAI barrier-confidence framework™ as a computational layer for quantifying assay resolution in an airway epithelial monolayer barrier OoC use case. The framework does not introduce or validate new biology. Instead, it asks a narrower question: given a defined assay, replicate design, target effect size, and uncertainty structure, is the target effect predicted to exceed the assay’s minimum detectable shift under a prespecified decision rule? Consistent with fit-for-purpose NAM principles [9,11], the context of use evaluated here is assay-readiness and decision support for laminar, low-Reynolds-number airway epithelial monolayer barrier OoC systems with documented relationships among flow, wall shear stress, and permeability-related readouts. The primary objective is to determine whether a predefined permeability-related effect can be resolved across reference and standardized workflow conditions that differ in modeled technical variability. The analysis is deliberately limited to in silico assay readiness and does not claim wet-lab validation, platform qualification, regulatory acceptance, clinical predictivity, or universal applicability of the modeled 10% target.

## 2 Methods

### 2.1 Barrier-confidence framework

The framework represents an observed assay readout as the combination of a biological signal and multiple uncertainty components, including manufacturing- and pump-driven variability, biological fluctuation, a run-level systemic component, and independent well-level noise (Figure 1). A reference workflow and a standardized workflow are assigned distinct technical-noise inputs while preserving the same biological target effect. Monte Carlo propagation is then used to estimate the distribution of assay resolution, expressed as the minimum detectable shift (MDS), and the target signal is evaluated against a fixed detectability rule. In the full framework, experimental data can subsequently be used to recalibrate uncertainty terms without changing the prespecified decision rule. That prospective calibration step has not yet been performed in the present simulation-only study.

**Figure 1.**
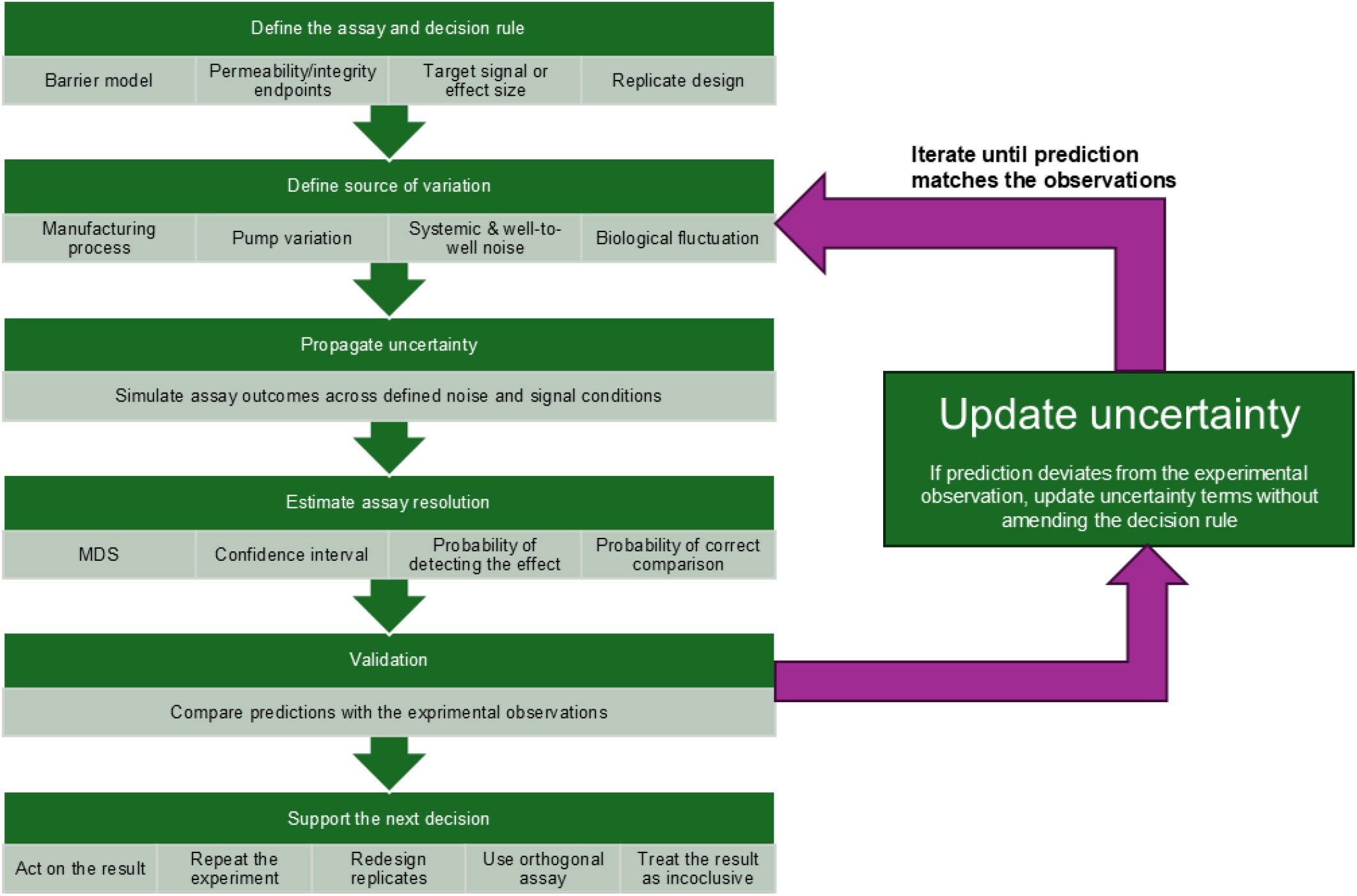
UstarFlowAI in silico barrier-confidence framework™. The computational layer sits on top of an existing barrier OoC workflow without changing the underlying biology. The full framework includes prospective comparison with experimental observations and, when required, recalibration of uncertainty terms while preserving the prespecified decision rule. The calibration loop shown here is proposed for future wet-lab validation and was not executed in the current simulation-only analysis.

### 2.2 Simulated variability domains

Monte Carlo simulation is designed to evaluate assay resolution across a range of nominal biological variability levels. Nominal biological coefficient of variation (CV) values, denoted *b*, ranged from 5% to 30% in 50 equally spaced increments. The CV is treated as an input parameter representing relative uncertainty in permeability for a single well and is not computed from the raw data. Biological CV, *B*_i_for a well *i* was modeled as an additive perturbation of the nominal biological CV:

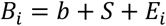

where:

- *b* is the nominal biological CV;
- *S* is the systemic component of the perturbation
- *E*_i_ is independent well-to-well component of the perturbation

The standard deviation of the perturbation is set to scale linearly with the nominal CV:

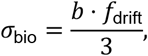

where biological drift factor *f*_drift_ = 0.20 for the best-case scenario, corresponding to a ±20% envelope around the nominal biological CV at three standard deviation levels, assuming normal distribution. The division by 3 implements a 3-sigma normalization, so that approximately 99.7% of wells have biological CV values within *b*⋅ (1 ± *f*_drift_), i.e., a ±20% envelope around the nominal CV. Similarly, the worst-case biological fluctuation envelope of ±30% is considered. The standard-deviation formulation for the perturbation is defined based on the assumption that the assay variability is fundamentally relative: a given percentage drift in assay performance (e.g., due to pipetting imprecision, flow instability etc.) produces larger absolute fluctuations in CV when the baseline CV, *b* is higher.

A mixed-noise structure comprising a correlated systemic component *S* and an independent well-level component *E* is used. For each Monte Carlo iteration, the systemic component was sampled once per simulated assay run:

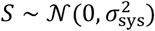

In the current implementation, the same systemic realization is shared across all wells in an assay run, including both control and treatment groups. The systemic component is therefore fully correlated across the two arms within an iteration.

The independent component was sampled separately for each well:

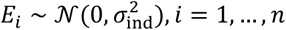

where:

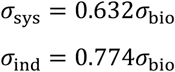

The coefficients approximate a variance allocation of 40% correlated systemic variation and 60% independent well-level variation for the worst-case envelope:

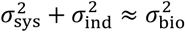

Similarly, coefficients approximating a variance allocation of 25% correlated systemic variation and 75% independent well-level variation were used for the best-case envelope:

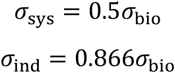

Two workflow conditions with distinct technical-noise inputs were modeled:

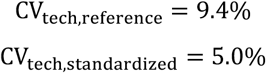

These fixed technical CV values represent model inputs for the reference planar barrier OoC and standardized barrier OoC workflows respectively. Complete set of parameter values used in the simulation are shown in Table 1. The physical design and computational methods used to derive the technical noise conditions are proprietary and are not disclosed in this manuscript. The total well-level noise for well *i*:

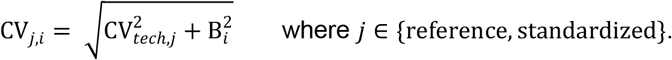

**Table 1.** Simulation parameters.

| Parameter domain | Current parameterization | Purpose |
| --- | --- | --- |
| Manufacturing and pump operating variability | Normal 3-sigma injection-molding and pump-variation assumptions are applied to both workflows. The derived technical-noise input is 9.4% CV for the reference workflow and 5.0% CV for the standardized workflow. Detailed tolerances and the proprietary method used to derive these values are not disclosed. | Represents fabrication- and pump-driven variability that can affect flow within the device. |
| Biological fluctuation | Scenario envelopes of $\pm 20\%$ (best case) and $\pm 30\%$ (worst case), modeled under a normal 3-sigma assumption, are used to stress-test plausible run-to-run and batch-to-batch variability in airway barrier OoC assays. | Represents biological fluctuation in the barrier response without claiming a specific cell line or protocol. |
| Systemic-to-well-level noise allocation | 25:75 in the best-case scenario and 40:60 in the worst-case scenario. | Stress-tests alternative allocations between correlated run-level and independent well-level variability. |
| Replicate number | $n = 48$ replicate wells per group (48 control and 48 treatment wells) for each workflow condition. | Evaluates whether a small target effect can be resolved under a high-replicate design. |
| Target effect | 10% permeability-related shift. | Illustrative treatment-response target used for assay-readiness classification. |
| Operating point | 17% nominal total assay CV. | Operating point used for the Figures 4 and 5 decision-boundary comparison. |

For each nominal biological CV value, 10,000 Monte Carlo iterations were performed. In each iteration, one systemic biological shift *S* and *n* = 48 independent well-level terms *E*_i_ were sampled for a simulated group. The total CV of each simulated well was calculated for both the reference and standardized technical-noise workflows. Nominal total CV across replicate wells was then calculated as:

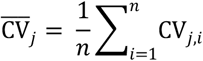

The arithmetic mean is used because each well-level CV_*j,i*_ is already a dimensionless, relative measure of uncertainty. Averaging CV_*j,i*_ provides a straightforward, interpretable summary of the typical assay-level CV across the simulated plate. This arithmetic summary is a model convention and is distinct from variance propagation across independent measurements. The current analysis uses *n* = 48 replicate wells per group, corresponding to a balanced two-arm design with 48 control wells and 48 treatment wells. A prespecified permeability-related shift of 10% was used as an illustrative assay-readiness target.

### 2.3 Assay-confidence metrics

The barrier-confidence framework uses two related quantities. First, the minimum detectable shift (MDS) estimates the smallest relative treatment-control difference expected to be detectable at a specified replicate density, significance level, power, and modeled total variability. Second, a conservative detectability rule compares the predefined target effect with the Monte Carlo distribution of MDS values at the selected operating point. A workflow is classified as PASS only when the target effect exceeds the upper bound of the 95% MDS interval; otherwise, it is classified as FAIL.

The MDS is estimated by a normal approximation formula for a two-sided comparison of two independent groups of equal size:

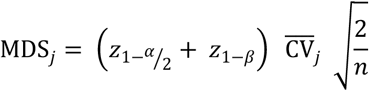

where:

- *α* = 0.05 is the two-sided type I error rate;
- 1 - *β* = 0.80 is the target statistical power;
- *n* = 48 is the number of replicate wells per group;
- 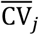 is the simulated mean total CV for workflow *j*.

Using the normal distribution approximation:

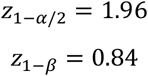

therefore:

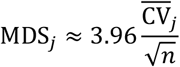

MDS is expressed as a percentage shift relative to the baseline assay readout. It represents the smallest relative treatment-control difference expected to be detectable with 80% power at a two-sided significance level of 0.05 under the stated assumptions of independent groups, equal variance, and approximately normal outcome distributions. Equal variability was assumed for control and treatment groups in the analytic MDS calculation. The simulation, however, includes a systemic component shared across both arms within each iteration. Because a common run-level component can contribute covariance to the treatment-control comparison, the closed-form MDS expression does not explicitly reproduce the full covariance structure of the simulation. In the present implementation, this simplification is used as a conservative assay-readiness approximation; its calibration against empirical data remains part of the prospective validation plan. For each nominal total-noise condition and respective workflow, Monte Carlo uncertainty ribbons were calculated from the 2.5th and 97.5th percentiles of the 10,000 simulated MDS values:

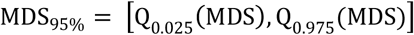

The nominal total CV of the reference condition was used as the x-axis variable to visualize MDS as a function of nominal total assay variability. For each operating point, Monte Carlo uncertainty ribbons summarize the distribution of predicted MDS values for reference and standardized workflows.

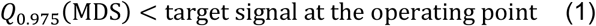

The decision rule in Equation (1) is intentionally conservative: the predefined target effect must remain above the upper 97.5th percentile of the simulated MDS distribution. If the target overlaps the MDS interval, or lies below its upper bound, the configuration is classified as FAIL under the specified assumptions. The lower 2.5th percentile is reported as part of the 95% interval but is not used for the PASS/FAIL classification. This rule is an assay-readiness criterion; it is not a direct estimate of experimental sensitivity, specificity, false-positive rate, false-negative rate, or clinical predictivity. Those properties require prospective empirical validation.

### 2.4 Simulation scenarios

Best- and worst-case simulation scenarios were evaluated. In both scenarios, assay resolution was compared between reference and standardized workflow conditions using n = 48 replicate wells per group across the modeled range of nominal biological variability (5%-30% CV). For each biological-noise level, 10,000 Monte Carlo iterations were performed. A predefined 10% permeability-related target effect was evaluated at a nominal total assay-noise operating point of 17% CV. The best- and worst-case scenarios used the lower and upper biological-fluctuation envelopes and the corresponding systemic-to-well-level noise allocations shown in Table 2.

**Table 2.** Best- and worst-case simulation scenarios.

| Scenario | Replicates per group | Manufacturing and pump assumptions | Operating point (nominal total CV) | Biological fluctuation envelope | Systemic:well-level allocation |
| --- | --- | --- | --- | --- | --- |
| Best case | 48 | Normal 3-sigma injection-molding and pump-variation assumptions | 17% CV | $\pm 20\%$ | 25:75 |
| Worst case | | | | $\pm 30\%$ | 40:60 |

All simulations were implemented in Python 3.12 using standard numerical libraries. The stochastic distributions, parameter ranges, Monte Carlo iteration count, MDS definition, and uncertainty-interval construction used in the present analysis are described in Sections 2.2–2.3 and Tables 1-2. The detailed physical-design specifications and proprietary computational methods used to derive the workflow-specific technical-noise inputs are not disclosed in this manuscript.

## 3. Results

### 3.1 Modeled reduction in technical variability

The shaded ribbons in Figures 2-5 represent the 95% Monte Carlo intervals of predicted MDS values across the defined uncertainty structure. The lower edge of a ribbon corresponds to lower-noise realizations and therefore better predicted assay resolution, whereas the upper edge corresponds to higher-noise realizations and poorer predicted resolution. The dashed line represents the mean MDS. These ribbons are simulation-derived uncertainty ranges and should not be interpreted as empirically measured day-to-day assay performance.

**Figure 2.**
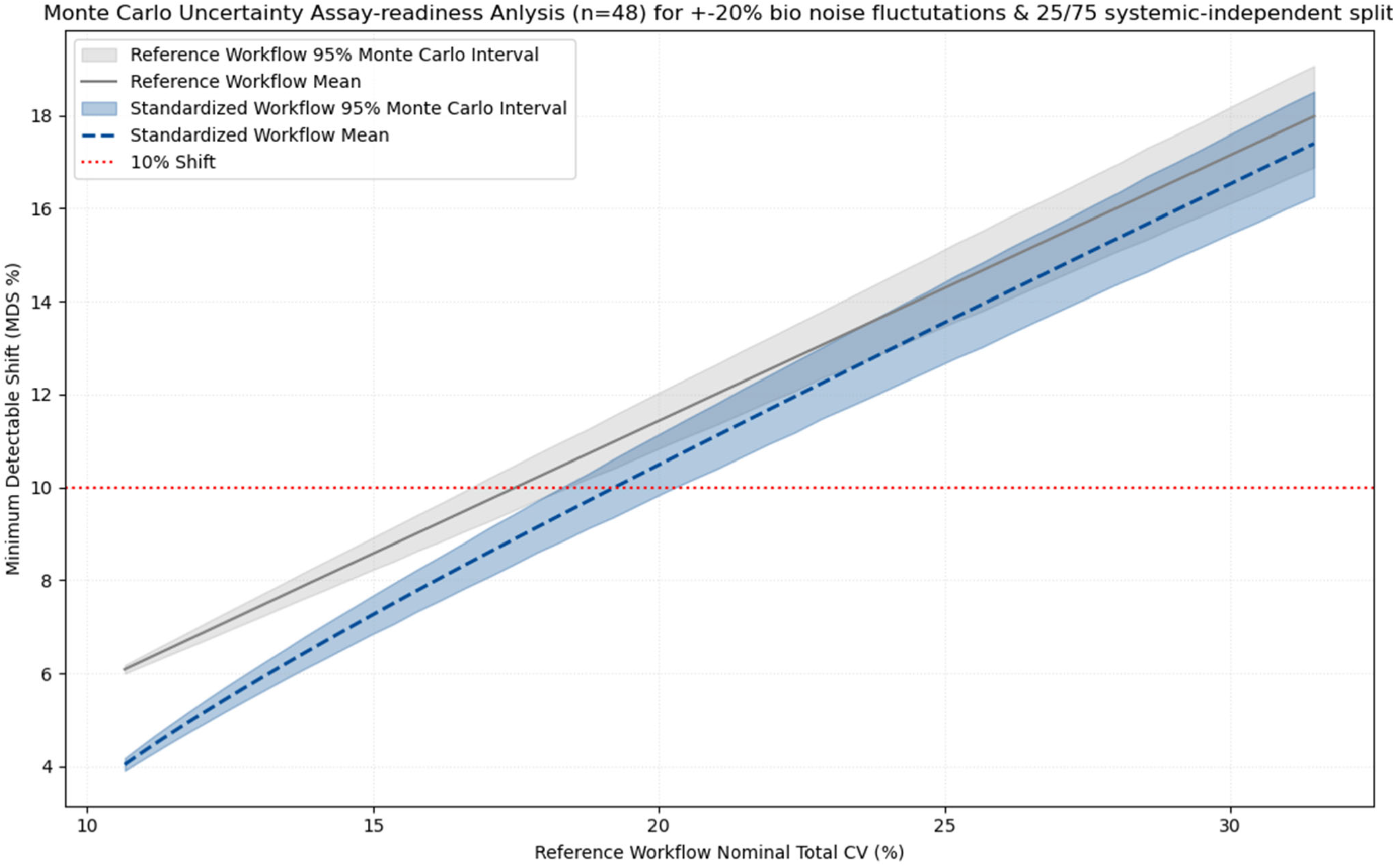
Illustrative in silico assay-readiness analysis for the best-case scenario. Predicted minimum detectable shift (MDS) for an airway epithelial monolayer barrier assay across a range of nominal assay-noise conditions with n = 48 replicate wells per group. Shaded ribbons show 95% Monte Carlo intervals; dashed lines show mean MDS. The standardized workflow is parameterized with lower technical noise than the reference workflow.

The reference and standardized workflows were parameterized with technical-noise inputs of 9.4% CV and 5.0% CV, respectively, corresponding to an approximately 46.8% reduction in the modeled technical-noise component (Figures 2 and 3). Because these values are inputs derived from proprietary manufacturing- and flow-related calculations, the present analysis demonstrates the consequence of that assumed reduction for assay resolution rather than experimentally establishing the reduction itself.

**Figure 3.**
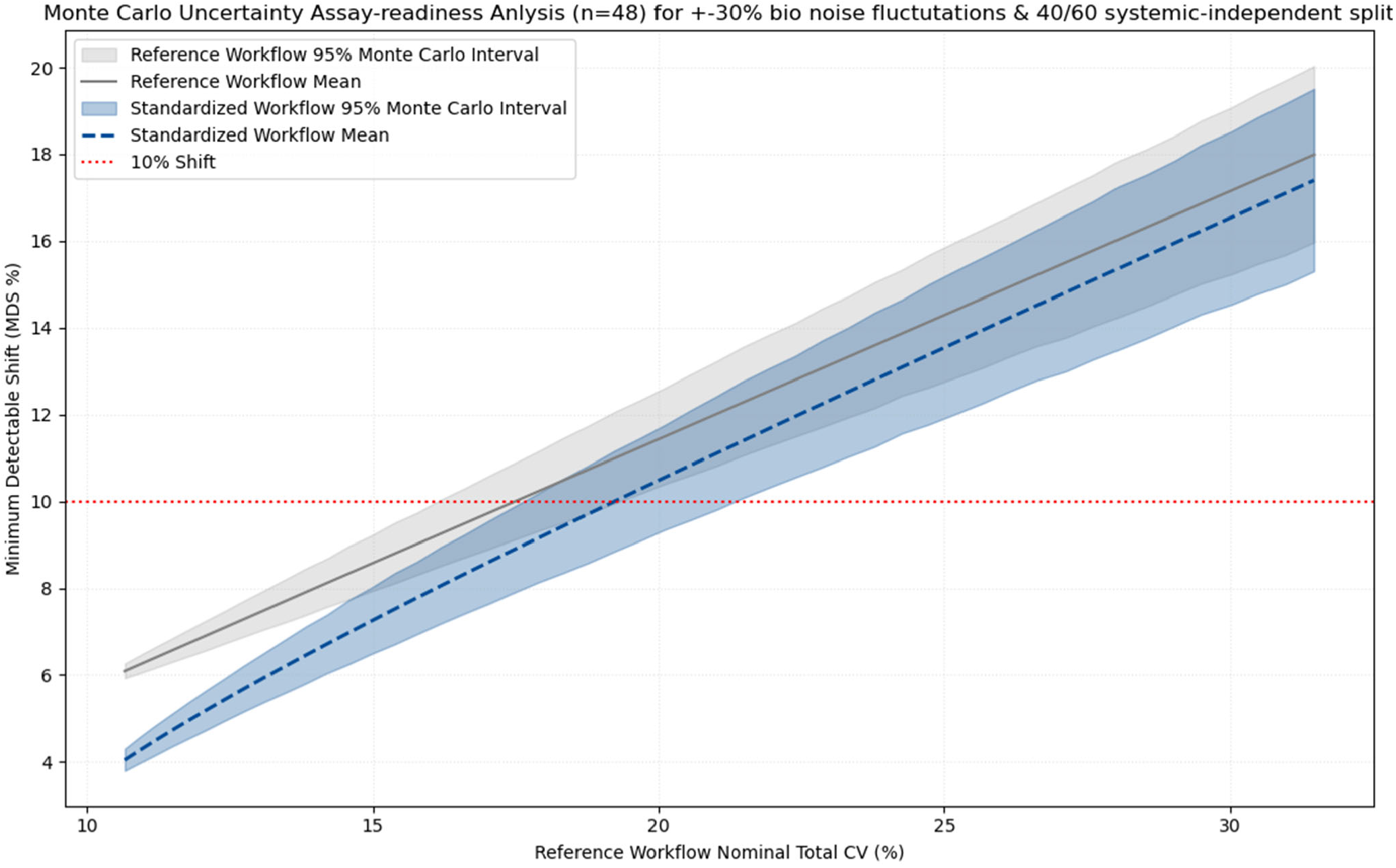
Illustrative in silico assay-readiness analysis for the worst-case scenario. Predicted MDS for an airway epithelial monolayer barrier assay across a range of nominal assay-noise conditions with n = 48 replicate wells per group. Shaded ribbons show 95% Monte Carlo intervals; dashed lines show mean MDS. The standardized workflow remains lower than the reference workflow across the modeled range, although biological uncertainty widens the intervals.

### 3.2 Improvement in modeled assay resolution

Across the evaluated range of nominal total assay noise, the standardized workflow produced a lower predicted MDS than the reference workflow (Figures 2 and 3). The magnitude of the difference depended on the relative contribution of technical and biological variability: when biological variability dominated the total noise, reducing the technical component produced a smaller absolute improvement in MDS; when technical variability contributed more strongly, the separation between workflows increased. All MDS values reported here are simulation outputs under the stated assumptions and are not experimentally measured assay-sensitivity estimates.

### 3.3 Detectability of a 10% permeability shift

Figures 4 and 5 show the decision-boundary analysis at the selected operating point of 17% nominal total CV. In both the best- and worst-case scenarios, the predefined 10% permeability shift remained above the upper MDS bound for the standardized workflow. The standardized workflow therefore met the prespecified PASS criterion in both scenarios.

**Figure 4.**
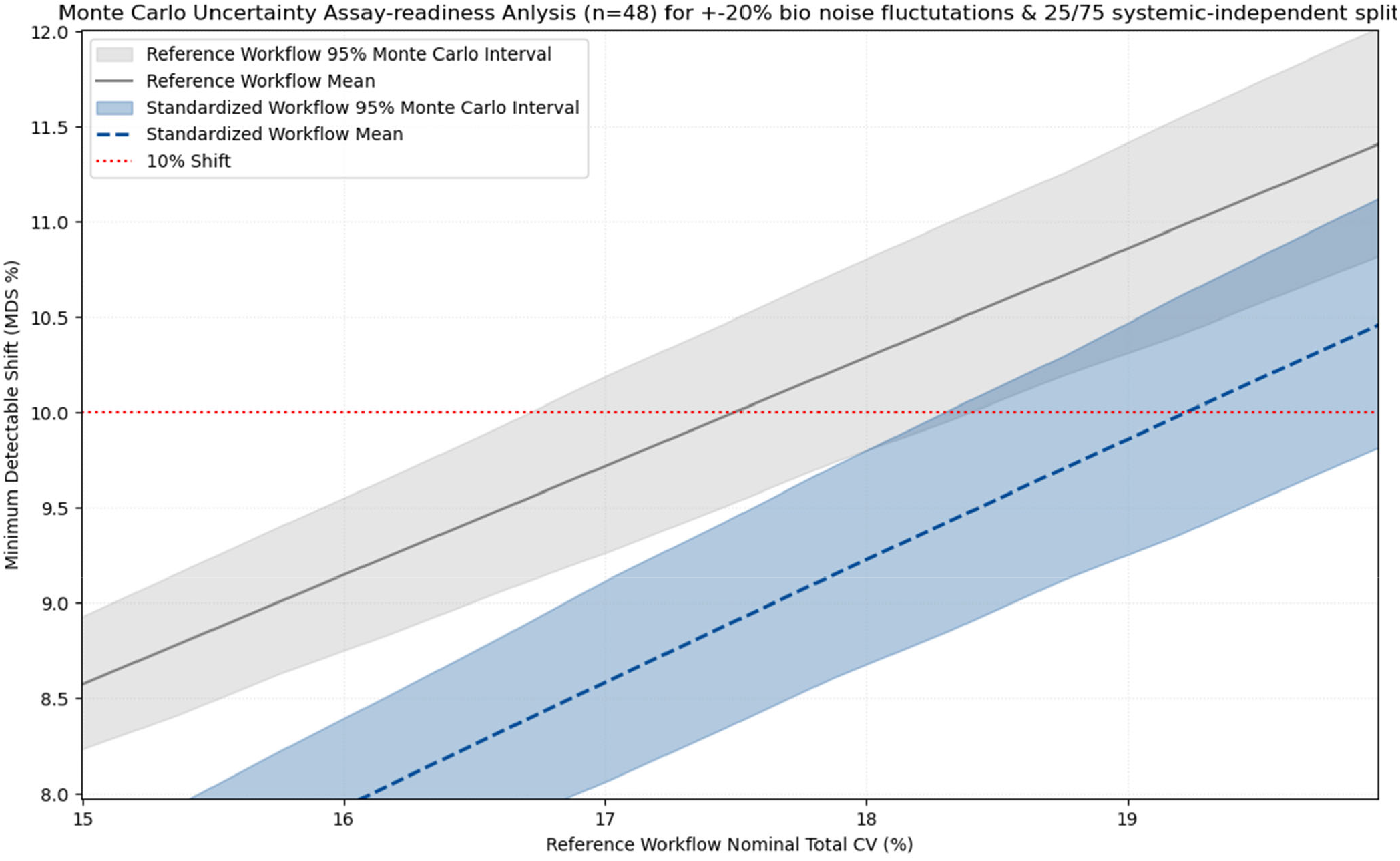
Illustrative in silico decision-boundary analysis for the best-case scenario. At the selected operating point of 17% nominal total CV and n = 48 replicate wells per group, the predefined 10% permeability shift remains above the upper MDS bound for the standardized workflow but overlaps the MDS interval for the reference workflow. Under the prespecified conservative rule, the standardized workflow is classified PASS and the reference workflow FAIL.

**Figure 5.**
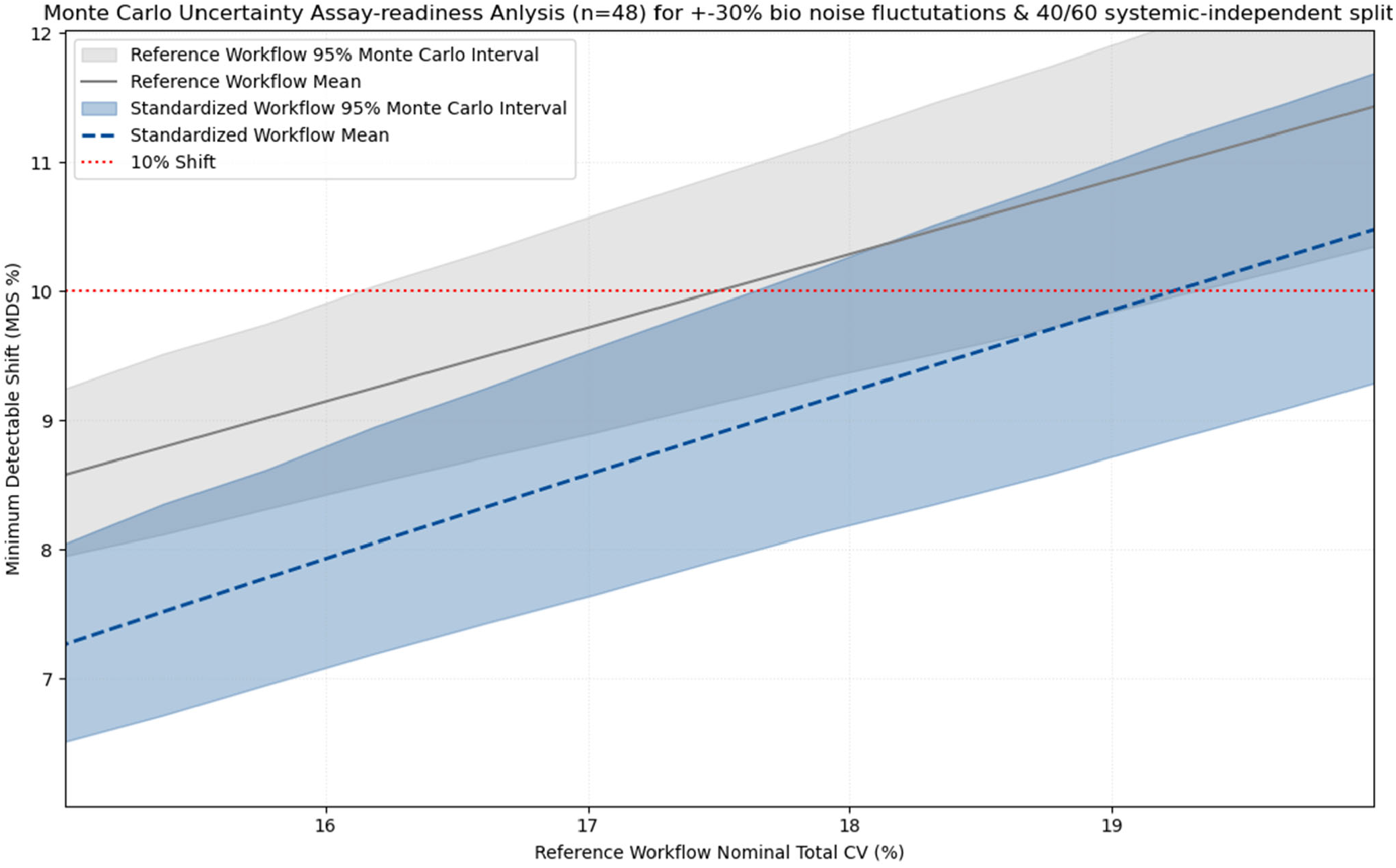
Illustrative in silico decision-boundary analysis for the worst-case scenario. At the same 17% nominal total-CV operating point and n = 48 replicate wells per group, the predefined 10% permeability shift remains above the upper MDS bound for the standardized workflow but overlaps the MDS interval for the reference workflow. The standardized workflow therefore remains PASS and the reference workflow FAIL under the prespecified rule.

For the reference workflow, the 10% target overlapped the MDS uncertainty interval at the same operating point in both scenarios. Although the mean MDS could remain below the target in some modeled conditions, the upper uncertainty bound did not. Under the conservative rule in Equation (1), the reference workflow was therefore classified as FAIL in both scenarios. This analysis does not estimate the empirical probability of detection; it identifies whether the target satisfies the prespecified uncertainty-bound criterion.

Together, these simulations show that, under the current parameterization and high-replicate design, reducing the modeled technical-noise component can shift a 10% target from an unresolved to a resolved region of the decision space. The result is conditional on the modeled device, uncertainty structure, n = 48 replicate wells per group, and 17% nominal total-CV operating point. It should not be generalized to other barrier OoC systems or interpreted as evidence that a 10% permeability change is inherently meaningful for a particular drug-development decision.

**Table 3.** Detectability stress-test results.

| Simulation | Replicates per group | Workflow | Endpoint | Result | Interpretation |
| --- | --- | --- | --- | --- | --- |
| Detectability stress test | 48 | Standardized | 10% permeability shift | PASS | Target exceeds the upper 95% MDS bound at the selected operating point. |
|  |  | Reference |  | FAIL | Target overlaps the 95% MDS interval at the selected operating point. |

## 4. Prospective Experimental Validation Strategy

Prospective wet-lab testing is required to determine whether the simulation-derived confidence metrics predict real assay behavior. A focused NAMina barrier-OoC study should therefore be designed prospectively, with the decision rule and target effect specified before outcome data are examined. Calibration and confirmatory experiments should be separated where feasible to reduce the risk of circular validation.

In a calibration phase, the study would use a single airway or other barrier OoC system with a stable, quantifiable permeability or TEER readout and a defined flow or wall-shear-stress range. Controlled device, flow, or pump perturbations would be introduced across the modeled technical-variability envelope without changing the underlying biological question. A modest treatment or perturbation effect would be prespecified for the selected assay; the current 10% target should be retained only if it is scientifically justified for that use case. Calibration data could then be used to update uncertainty terms while preserving the prespecified decision rule.

In a separate confirmatory phase, the expected MDS and PASS/FAIL prediction would be calculated before unblinding experimental outcomes. Predicted detectability would then be compared with observed treatment-control effect estimates and confidence intervals. After a sufficient number of independent runs, concordance between prediction and observation, false-positive and false-negative behavior, and sensitivity to replicate number and uncertainty allocation could be evaluated.

Transferability should be assessed by repeating the confirmatory analysis across at least one independent batch, operator, or laboratory condition. This design would test whether the calibrated uncertainty model remains informative outside the conditions used for calibration and would provide the empirical basis needed to judge whether the framework is fit for the intended context of use.

## 5. Discussion

The principal contribution of the barrier-confidence framework is methodological: it shifts the question from generic platform reproducibility to effect-specific assay resolution. A barrier model may be biologically relevant yet still be insufficiently resolved for a particular decision if the expected effect is smaller than the uncertainty surrounding the assay. Conversely, an assay does not need to eliminate all variability to be useful; it needs to characterize variability well enough to determine whether the effect of interest is distinguishable under the intended conditions.

The framework complements rather than replaces conventional power analysis. Standard power calculations typically ask how many replicates are required to detect an assumed effect at an assumed variance. Here, the additional objective is to decompose uncertainty into biologically and technically interpretable components and to ask whether changing a workflow-specific technical component materially shifts the decision boundary. This framing is consistent with fit-for-purpose MPS quality-management principles and current NAM guidance, which emphasize context of use, reproducibility, transparent performance characteristics, and decision relevance [9,11].

The present simulations illustrate this principle. At the selected 17% nominal total-CV operating point and n = 48 replicate wells per group, the standardized workflow met the conservative detectability criterion for a 10% target in both best- and worst-case scenarios, whereas the reference workflow did not. This difference is driven by the workflow-specific technical-noise inputs and should not be interpreted as experimental proof of superiority. The modeled values are conditional on the assumptions used to derive and propagate uncertainty.

The treatment-control contrast also requires explicit statistical qualification. In the current simulation, the systemic realization is shared across both arms within an assay run. A common run-level component can therefore contribute covariance to the contrast, whereas the closed-form MDS calculation uses a simplified independent-groups approximation based on marginal CV. The approximation does not explicitly model that covariance structure. In the present implementation it is intended to provide a conservative assay-readiness screen, but the degree of conservatism has not yet been quantified empirically. Prospective calibration should therefore test not only PASS/FAIL concordance but also whether predicted MDS values are appropriately calibrated across operating conditions.

A second design principle is separation of the biological model from the confidence layer. Biological relevance remains determined by the OoC system, cell source, culture conditions, endpoint selection, and experimental protocol. The computational layer instead evaluates whether the resulting measurement is sufficiently resolved for a predefined decision. This modular framing could, after empirical validation, allow the same uncertainty-based logic to be evaluated across different OoC technologies without transferring ownership of platform-specific biology or proprietary protocols. Evidence of framework utility would therefore require prospective agreement between predicted and observed detectability, stability of the calibration across independent runs, and transparent definition of the context in which the prediction is intended to support a decision.

## 6. Limitations

- The current evidence is simulation-based and has not yet been prospectively validated against wet-lab data.
- Workflow-specific technical CV values are model inputs derived using proprietary manufacturing-and flow-related methods; their empirical accuracy remains to be demonstrated.
- The analytic MDS approximation assumes independent groups, equal variance, and approximately normal outcomes. The simulation includes a shared systemic component, so the closed-form equation does not explicitly represent the full covariance structure of the two-arm assay.
- The 10% permeability target is illustrative and has not been established as a universally biologically or clinically meaningful effect or as an established regulatory threshold.
- The present results are specific to the modeled barrier/permeability use case and should not be extrapolated to other OoC endpoints, devices, replicate structures, or contexts of use without separate evaluation.
- The MDS calculation assumes equal variance in control and treatment groups. If an intervention changes the variability of the barrier readout as well as its mean, the resulting detection threshold may differ from the present estimate.

## 7. Conclusions

The UstarFlowAI barrier-confidence framework™ provides a quantitative way to express whether a predefined effect is resolvable under specified assay uncertainty. In this in silico proof of concept, a standardized workflow parameterized with lower technical variability produced lower MDS values than the reference workflow and met the prespecified criterion for resolving a 10% permeability shift at n = 48 replicate wells per group and a 17% nominal total-CV operating point across both modeled scenarios. The reference workflow did not meet that conservative criterion. These findings establish a testable computational hypothesis rather than experimental validation. The next critical step is prospective wet-lab evaluation of whether simulation-derived confidence metrics predict real assay detectability and remain calibrated across independent experimental conditions.

## Author Contributions

Paola Dama: conceptualization of the translational context of use; experimental-validation strategy; writing - original draft; writing - review and editing. Shrawasti Sahare: conceptualization of the UstarFlowAI barrier-confidence framework™; simulation design and analysis; technical interpretation; visualization; writing - review and editing. Both authors contributed to study-design refinement and manuscript development.

## Data and Code Availability

No new experimental datasets were generated for this study. The simulation parameter values, statistical assumptions, and derived outputs required to interpret the analyses are reported in the manuscript. Detailed manufacturing tolerances, pump-performance specifications, proprietary methods used to derive workflow-specific technical-noise inputs, and source code are not publicly available because they constitute proprietary methods of UstarFlowAI. Non-proprietary derived simulation outputs may be made available by the co-corresponding authors upon reasonable request.

## References

1. Huh D, Matthews BD, Mammoto A, Montoya-Zavala M, Hsin HY, Ingber DE. Reconstituting organ-level lung functions on a chip. Science. 2010;328(5986):1662–1668. doi:10.1126/science.1188302.

2. Lagowala DA, Kwon S, Sidhaye VK, Kim DH. Human microphysiological models of airway and alveolar epithelia. Am J Physiol Lung Cell Mol Physiol. 2021;321(6):L1072–L1088. doi:10.1152/ajplung.00103.2021.

3. Arik YB, van der Helm MW, Odijk M, Segerink LI, Passier R, van den Berg A, van der Meer AD. Barriers-on-chips: Measurement of barrier function of tissues in organs-on-chips. Biomicrofluidics. 2018;12(4):042218. doi:10.1063/1.5023041.

4. Blume C, Reale R, Held M, Millar TM, Collins JE, Davies DE, Morgan H, Swindle EJ. Temporal monitoring of differentiated human airway epithelial cells using microfluidics. PLoS One. 2015;10(10):e0139872. doi:10.1371/journal.pone.0139872.

5. Sidhaye VK, Schweitzer KS, Caterina MJ, Shimoda L, King LS. Shear stress regulates aquaporin-5 and airway epithelial barrier function. Proc Natl Acad Sci U S A. 2008;105(9):3345–3350. doi:10.1073/pnas.0712287105.

6. Sidhaye VK, Chau E, Breysse PN, King LS. Septin-2 mediates airway epithelial barrier function in physiologic and pathologic conditions. Am J Respir Cell Mol Biol. 2011;45(1):120–126. doi:10.1165/rcmb.2010-0235OC.

7. Holzreuter MA, Segerink LI. Innovative electrode and chip designs for transendothelial electrical resistance measurements in organs-on-chips. Lab Chip. 2024;24:1121–1134. doi:10.1039/D3LC00901G.

8. Emulate Bio. Protocol for Emulate Organ-Chips: Barrier Function Analysis. Document EP187, version 1.0. 2019.

9. Pamies D, Ekert J, Zurich MG, et al. Recommendations on fit-for-purpose criteria to establish quality management for microphysiological systems and for monitoring their reproducibility. Stem Cell Reports. 2024;19(5):604–617. doi:10.1016/j.stemcr.2024.03.009.

10. Braakhuis HM, Gremmer ER, Bannuscher A, et al. Transferability and reproducibility of exposed air-liquid interface co-culture lung models. NanoImpact. 2023;31:100466. doi:10.1016/j.impact.2023.100466.

11. U.S. Food and Drug Administration, Center for Drug Evaluation and Research. General Considerations for the Use of New Approach Methodologies in Drug Development: Draft Guidance for Industry. March 2026.

